# Identification of a human adult cardiac stem cell population with neural crest origin

**DOI:** 10.1101/590679

**Authors:** Anna Höving, Madlen Merten, Kazuko Elena Schmidt, Isabel Faust, Lucia Mercedes Ruiz-Perera, Henning Hachmeister, Sebastian-Patrick Sommer, Buntaro Fujita, Thomas Pühler, Thomas Huser, Johannes Greiner, Barbara Kaltschmidt, Jan Gummert, Cornelius Knabbe, Christian Kaltschmidt

**Author notes:** these authors contributed equally to this work (Anna Höving, Madlen Merten, Barbara Kaltschmidt, Cornelius Knabbe, Christian Kaltschmidt).

## Abstract

Cardiovascular diseases are the major cause of death worldwide, emphasizing the necessity to better understand adult human cardiac cell biology and development. Although the adult heart was considered as a terminally differentiated organ, rare populations of cardiac stem cells (CSCs) have been described so far, with their developmental origin and endogenous function still being a matter of debate.

Here, we identified a Nestin^+^/S100^+^/CD105^+^/Sca1^+^/cKit^-^ population of CSCs in the myocardium of the adult human heart auricle. Isolated cells showed expression of characteristic neural crest-derived stem cell (NCSC) markers and kept their genetic stability during cultivation *in vitro*. Cultivated hCSCs efficiently gave rise to functional, beating cardiomyocytes, osteoblasts, adipocytes and neurons. Global transcriptome analysis via RNAseq showed a high similarity between the expression profiles of Nestin^+^/S100^+^/CD105^+^/Sca1^+^/cKit^-^ hCSCs and adult human NCSCs from the nasal cavity (inferior turbinate stem cells, ITSCs). In detail, 88.1 % of all genes were significantly expressed in both stem cell populations particularly including common NCSC-markers. Based on these observations, we suggest a similar developmental origin of both stem cell populations.

In summary, we identified a human adult cardiac stem cell population with neural crest-origin, which may also contribute to endogenous cardiac tissue homeostasis and tissue repair *in vivo.*

## INTRODUCTION

Cardiovascular diseases are the major cause of death worldwide ^1^ and main clinical treatments are still palliative, emphasizing the necessity to establish suitable *in vitro* models to better understand adult human cardiac cell biology. Although the adult heart was long time considered as a terminally differentiated organ, several populations of adult cardiac stem cells (CSCs) have been found recently in the mouse and human heart^2-4^. These cells are mainly able to differentiate into cardiomyocytes but also into smooth muscle cells as well as endothelial cells *in vitro* ^2,5,6^. *Nevertheless, the in vivo* contribution of CSCs to tissue repair remains elusive, partly because of the heterogeneity of the CSC populations described so far^7,8^. While adult cardiac stem cells were initially described and isolated according to their expression of the stem cell factor receptor kinase cKit ^2,9,10^, Isl1^+^ and Sca1^+^ cardiac stem cell populations have also been described during the last years ^11-13^. These observations led to the assumption that the so far characterized variation of adult stem cell populations may depend on their developmental origin. The neural crest was initially described by Wilhelm His in the development of the chick embryo as the intermediate chord appearing between the neural chord and the future ectoderm ^14^ and is often considered as a fourth germ layer. Among the broad range of adult human stem cells, neural crest-derived stem cells (NCSCs) show an extraordinary high differentiation potential *in vivo* and *in vitro* and are therefore considered as extended multipotent and promising candidates for clinical applications^15-17^. Although NCSCs can be found in various tissues of the adult human body^18-21^, a neural crest origin of CSCs has only been described in the murine heart^22,23^.

The present study aims to investigate the expression profile, differentiation potential and developmental origin of human CSCs (hCSCs) in more detail, particularly regarding a potential relation to the neural crest. We show that S100^+^/Nestin^+^ adult neural crest-derived stem cells can be found in the human left atrial appendage (LAA) and provide a method for isolation and culture of these cells. Further, these cells exhibit the characteristic expression profile of NCSCs and give rise to neuron-like cells, osteoblasts, adipocytes and functional cardiomyocytes *in vitro*. While previous characterizations of human cardiac stem cells focused on the presence of cell surface markers, global gene expression profiles have not been investigated so far. Here, RNAseq served to compare the whole transcriptome of hCSCs with inferior turbinate stem cells (ITSCs), a well characterized NCSC population from the human nose ^15,18,24^. Both adult stem cell populations showed high similarities in global gene expression particularly regarding neural crest markers like *NESTIN, SOX9, SNAIL, SLUG* and *TWIST*. Our study therefore presents a novel cKit^-^ stem cell population in the adult human heart with characteristics commonly associated with NCSC. Regarding the extraordinary high differentiation potential of NCSCs and the successful differentiation of NC-derived hCSCs into functional cardiomyocytes *in vitro*, Nestin^+^/cKit^-^ CSCs might serve as a promising source for therapeutic applications.

## METHODS

### Cell Isolation and Cultivation

Human left atrial appendage tissue was obtained from patients during routine surgery at Heart and Diabetes Centre NRW (Bad Oeynhausen, Germany) after informed written consent according to local and international guidelines (declaration of Helsinki). Isolation and further experimental procedures were ethically approved by the ethics commission of the Ruhr-University Bochum (Faculty of medicine, located in Bad Oeynhausen) (approval reference number eP-2016-148). After surgical removal, biopsies were cut into small pieces, and washed in PBS (Sigma Aldrich). For initial expansion, the tissue clumps were seeded in gelatin B-coated 10 cm-petri-dishes (Sarstedt AG & Co.) with human cardiac stem cell medium (hCSC-medium) consisting of DMEM/F-12 medium (Sigma Aldrich), basic fibroblast growth factor (bFGF, 5 ng/ml; Peprotech), epidermal growth factor (EGF, 10 ng/ml; Peprotech) and 10% fetal calf serum (VWR). After reaching confluence, tissue clumps were removed and passaging was performed by treatment with trypsin-EDTA (Sigma Aldrich). For further cultivation, cells were again seeded in gelatin B-coated T-25 cell culture flasks (Sarstedt AG & Co.) in hCSC-medium. For clonal analysis, cells were seeded in a 96-well plate (Sarstedt AG & Co.) at a density of 1 cell per well. Single cell dilution was verified by microscopy and medium was changed all two to three days. Sphere forming capacity was tested in low-adhesion culture flasks (Greiner Bio-One) with stem cell medium^18^.

Human inferior turbinate stem cells were isolated and cultivated after informed written consent according to local and international guidelines (declaration of Helsinki) as previously described ^18,24^. Isolation and further experimental procedures were ethically approved by the ethics commission of the Ärztekammer Westfalen-Lippe and the medical faculty of the Westfälische Wilhelms-Universität (Münster, Germany) (approval reference number 2012-15-fS).

### Lentiviral transduction of hCSCs

HCSCs were transduced by lentivirus with the cFUG-W plasmid. Lentivirus production was carried out in HEK293 cells with packaging plasmid Δ8.91, VSV-G envelope plasmid and cFUG-W transfer vector by calcium-phosphate precipitation. Δ8.91 and VSV-G were gifts from David Baltimore^25^. Supernatant was harvested 48h after transfection and lentivirus was concentrated by ultracentrifugation (50000 x g, 4°C, 2h).

### Coculture of GFP-hCSCs and primary mouse cardiomyocytes

Primary mouse cardiomyocytes were isolated from newborn mice according to Streejt and colleagues ^26^. Prior to coculture, mouse cardiomyocytes were treated with 10 µg/ml Mitomycin C (Sigma Aldrich) according to manufacturer’s instructions. Coculture with hCSCs was carried out in DMEM-F12 with 5 % horse serum (Dianova).

### Immunohistochemistry and Immunocytochemistry

Cryosections of the heart auricle tissue or cultivated cells were fixed for 20 min using 4% paraformaldehyde (PFA), washed and permeabilized in PBS with TritonX-100 (tissue: 0.2%, cells: 0.02%, Applichem) and supplemented with 5% goat serum for 30 min. The applied primary antibodies were diluted in PBS as followed: rabbit anti-Nestin 1:200 (Millipore), mouse anti-S100B 1:500 (Sigma Aldrich), rabbit anti-Slug 1:100 (Cell-Signaling Technology), rabbit anti-p75 1:500 (Cell-Signaling Technology), mouse anti-β-III-tubulin 1:100 (Promega), rabbit anti-neurofilament-L 1:50 (Cell-Signaling Technology), mouse anti-α-actinin (Cell-Signaling), anti-vGlut (Millipore) and anti-Synaptophysin (Millipore). They were applied for 1 h (cells) or for 2 h (sections), both at RT. After three washing steps, secondary fluorochrome-conjugated antibodies (Alexa 555 anti-mouse or Alexa 488 anti-rabbit, Invitrogen, Life Technologies GmbH) were applied for 1 h at RT with a dilution ratio of 1:300. Nuclear staining was realized by incubation with 4,6-Diamidin-2-phenylindol (DAPI) (1 μg/ml, Applichem) in PBS for 15 min at RT. Finally, the samples were mounted with Mowiol (self-made). Imaging was performed using a confocal laser scanning microscope (CLSM 780, Carl Zeiss) and image processing was executed with ImageJ and CorelDRAW ^27^ (open source and Corel Corporation).

### Quantitative PCR

The RNA isolation was performed utilizing the NucleoSpin RNA Kit (Macherey Nagel) according to the manufacturer’s guidelines. The quality and concentration of the obtained RNA was quantified by a spectrophotometer (Thermo Fisher Scientific). For cDNA synthesis, the First Strand cDNA Synthesis Kit (Thermo Fisher Scientific) was applied in accordance to the manufacturer’s guidelines. qPCR was carried out using Perfecta SYBR green Supermix (quantaBio) following manufacturer’s instructions with primers for *NESTIN* (fwd: CAGCGTTGGAACAGAGGTTG, rev: GCTGGCACAGGTGTCTCAAG), *CMYC* (fwd: AGGAGACATGGTGAACCAGAGT, rev: AGCCTGCCTCTTTTCCACAGAAAC), *OCT4* (fwd: CGAAAGAGAAAGCGAACCAG, rev: GCCGGTTACAGAACCACACT), *SOX10* (fwd: AGGAGAAGGAGGTTGACTGT, rev: TCCTCAAAGCTACTCTCAGC), *SOX9* (fwd: CCTCTCGCTTCAGGTCAGC, rev: ACTCGCCACACTCCTCCTC), *SNAIL* (fwd: CCCAATCGGAAGCCTAACTA, rev: GGACAGAGTCCCAGATGAGC), *SLUG* (fwd: TCGGACCCACACATTACCTT, rev: TTGGAGCAGTTTTTGCACTG), *TWIST* (fwd: GTCCGCAGTCTTACGAGGAG, rev: CCAGCTTGAGGGTCTGAATC), *LYPD6* (fwd: GCTCCAGACATCTACTGCCC, rev: AGAAGTGCAGACCTTGTGGC), *PPARG* (fwd: GGATGCAAGGGTTTCTTCCG, rev: AACAGCTTCTCCTTCTCGGC), *ON* (fwd: AAACATGGCAAGGTGTGTGA, rev: TGCATGGTCCGATGTAGTC) and *GAPDH* (fwd: CATGAGAAGTATGACAACAGCCT, rev: AGTCCTTCCACGATACCAAAGT).

### Flow Cytometry

Cultivated hCSCs were harvested by centrifugation after treatment with trypsin and subsequently stained with PE-coupled anti-CD105, anti-CD117, anti Sca1 or anti-CD31 antibody (Miltenyi Biotec, Bergisch Gladbach, Germany) according to manufacturer’s guidelines. For isotype controls, hCSCs were stained with PE-coupled IgG1 control antibody or APC-coupled IgG1 control antibody. Analysis was done using Gallios Flow Cytometer (Beckmann Coulter Inc., Brea, CA, United States), while Kaluza Acquisition Software (Beckmann Coulter Inc.) served for subsequent data acquisition and statistical analysis.

For DNA content analysis, nuclei were stained with propidium iodide (PI). Briefly, cells were lysed in 70% ethanol for 5 minutes and after centrifugation a staining solution of PBS containing glucose (1 mg/ml) (Carl Roth GmbH) and propidium iodide (12 µl/ ml) (Partec GmbH) was applied.

### Differentiation into Osteoblasts

The osteogenic differentiation of hCSCs was induced by biochemical cues according to Greiner and coworkers ^24^. Briefly, cells were seeded in hCSC-Medium at a density of 3 × 10^3^ cells/cm^2^. After 48 h the medium was switched to an osteogenic induction medium supplemented with 100 nM dexamethasone (Sigma Aldrich), 0.05 mM L-ascorbic acid-2-phosphate (Sigma Aldrich) and 10 mM β-glycerophosphate (Sigma Aldrich). The medium was changed every two to three days. After 21 days differentiated cells were processed for RNA-Isolation or Alizarin red S staining as described below. For undifferentiated controls, cells were cultured in hCSC-medium as described above.

### Alizarin Red Staining

For detection of calcium deposits maturated cells were fixed for 20 min with 4% PFA and briefly washed with PBS followed by a thorough washing with ddH2O. Subsequently, a staining solution of 1% Alizarin red S (Waldeck) in ddH2O with a pH of 4.3 was applied for 5 min at RT followed by rinsing with ddH2O.

### Induced Neuronal Differentiation

Neuronal differentiation in the isolated cells was induced following the protocol described by Müller and colleagues^15^. Briefly, cells were seeded with a density of 2 × 10^5^ cells per 6-well in hCSC-Medium. After 48 h, neural differentiation was induced with a neuronal induction medium containing 1 μM dexamethasone (Sigma Aldrich), 2 μM insulin (Sigma Aldrich), 500 μM 3-isobutyl-1-methylxanthine (Sigma Aldrich) and 200 μM indomethacin (Sigma Aldrich). Cells were fed every 2-3 days by removing half of the medium and adding the same amount of fresh prewarmed medium. After 7 days of culture, maturation of the cells was induced by adding retinoic acid (0.5 mM) (Sigma Aldrich) and N2-supplement (1x) (Gibco) over two days. Afterwards, retinoic acid was removed while N2 was applied until neuronal maturation at day 28. As undifferentiated control, cells were cultured in hCSC-medium as described above. After 28 days, the protein expression was analyzed by immunocytochemical staining.

### Adipogenic Differentiation of hCSCs

For adipogenic differentiation, hCSCs were cultivated in DMEM (Sigma Aldrich) containing 10% FCS (Sigma Aldrich) and plated at a density of 4 × 10^3^ cells /cm^2^. After 48 h, 1 μM dexamethasone (Sigma Aldrich), 2 μM insulin (Sigma Aldrich), 500 μM 3-isobuthyl-1-methylxanthine (Sigma Aldrich) and 200 μM indomethacin (Sigma Aldrich) were added to the medium and cultivated for 72 h. Afterwards the medium was switched and cells were cultivated for 4 days in DMEM containing 10% FCS and 2 μM insulin (Sigma Aldrich) to induce adipogenic differentiation. These two media were alternatingly used and changed every 4 days for 3 weeks. Subsequently, cells were fixed in 4% PFA for 1 h and stained with Oil red O (Sigma Aldrich). As undifferentiated control, cells were cultured in hCSC-medium as described above.

### Cardiac differentiation of hCSCs

Cardiac differentiation of the isolated cells was induced following the protocol described by Smits and colleagues ^4^. Briefly, cells were seeded with a density of 10^5^ cells per 6-well in hCSC-Medium. After 24 h, differentiation was induced with a cardiac differentiation medium consisting of a 1:1-mixture of IMDM (Gibco, Thermo Fisher Scientific) and Ham’s F12 nutrient mixture with GlutaMAX-I (Gibco), containing 10% horse serum (Dianova), 1x MEM nonessential amino acids (Bio Whittaker) and 1x insulin-transferrin-selenium (Gibco). 5 µM 5-azacytidine was added in three consecutive days and differentiation medium was refreshed at day 4. Six days after start of the differentiation, ascorbic acid (Sigma Aldrich) was added every two days and 1 ng/ml transforming growth factor β (TGF-β) (Peprotech) was added twice weekly. Medium was refreshed every two to three days. After 28 days, the protein expression was analyzed by immunocytochemical staining for α-actinin as described above. As undifferentiated control, cells were cultured in hCSC-medium.

### RNA-seq and Bioinformatics

RNA of cultured cells was isolated as described above and stabilized with RNAstable (Biomatrica, San Diego) for transport at room temperature. RNA was sequenced by Novogene (Beijing, China) using the Illumina Hiseq4000 platform with a paired end 150 bp strategy. Clean reads were aligned to the reference genome homo sapiens (GRCh37/hg19) using TopHat v2.0.9 and HTSeq v0.6.1 was used to count the read number mapped for each gene. RNAseq data are accessible at NCBI Gene Expression Omnibus. Differential gene expression analysis between two groups was performed using the DESeq2 R package (2_1.6.3). Correlation was determined using the cor.test function in R. GO terms of similar expressed genes were determined using the PANTHER classification system ^28^ and analysis of KEGG pathway enrichment was performed using KOBAS 3.0 ^29,30^. Principal component analysis (PCA) was carried out using MATLAB from MathWorks and the build-in PCA function.

## RESULTS

### Identification of NCSC-marker expressing cells in human heart auricle tissue

To localize putative adult cardiac stem cell populations within their endogenous niche, human heart auricles (LAA = left atrial appendage) (Fig. 1 A) were obtained during routine heart surgery. Within this heart auricle tissue a typical morphology of three main layers was observable: The myocardium is cardiac muscle tissue, mainly cardiomyocytes that is surrounded by the epicardium to the outer surface and the endocardium to the inner surface of the heart (Fig. 1 B). Immunohistochemistry revealed the presence of cells positive for the neural-crest markers S100 and Nestin in the adult myocardium and not in the endocardium or epicardium (Fig. 1 C). After RNA Isolation from heart auricle tissue samples, qPCR verified the expression of the SC-marker *NESTIN* on mRNA-Level. Moreover, expression of additional NCSC-markers like *SOX9, SOX10, SLUG* and *TWIST* was detectable next to expression of pluripotency markers *CMYC* and *OCT4* (Fig. 1 D+E).

**Figure 1:**
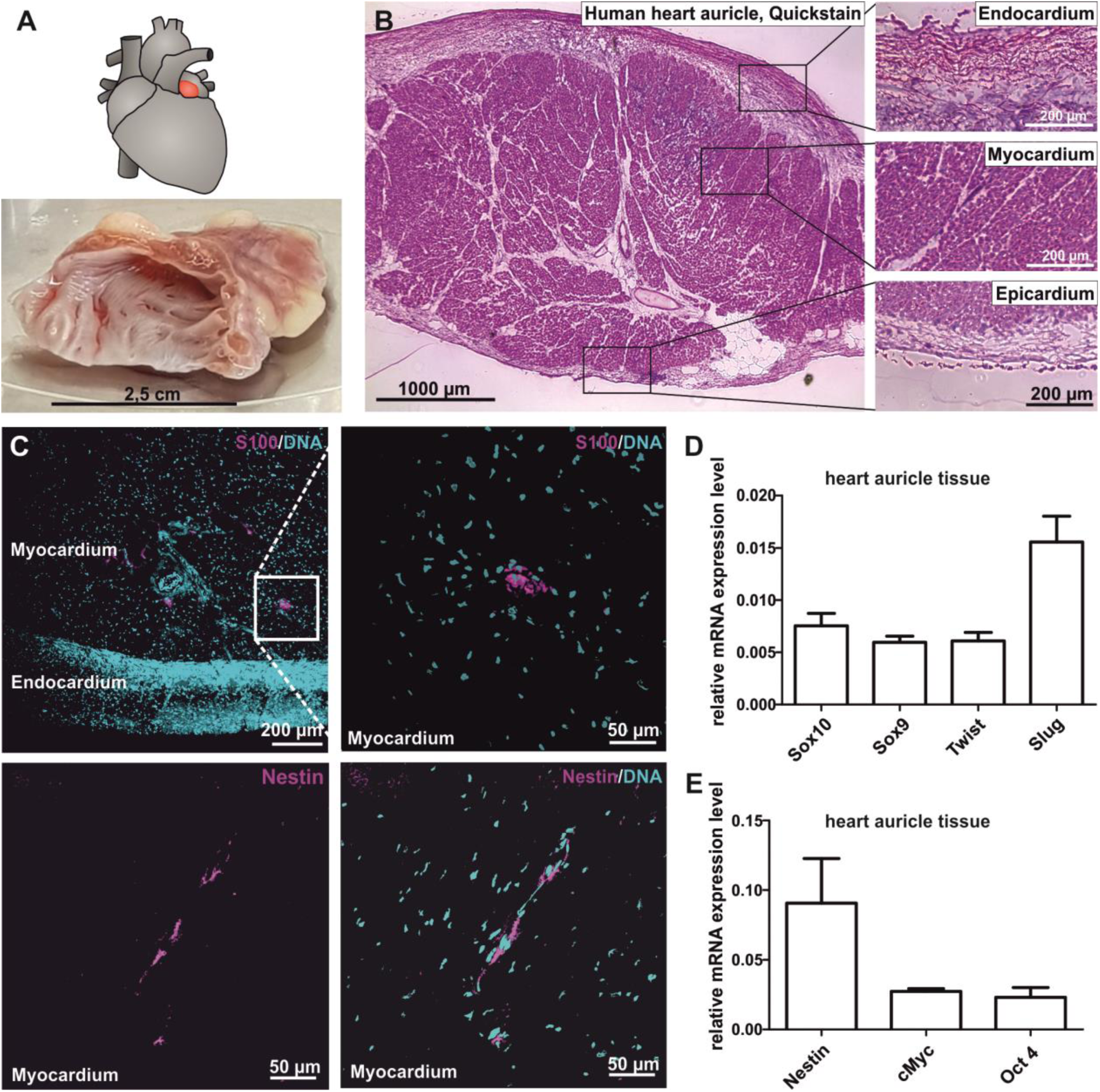
Human heart auricle tissue contains cells expressing markers characteristic for neural crest-derived stem cells. (A) Schematic view of the human heart, the heart auricle is indicated in red. A large amount of tissue can be obtained during surgery. (B) HE staining showing the characteristic tissue layers of endocardium, myocardium and epicardium. (C) Immunohistochemically staining revealed the presence of cells expressing Nestin and S100B in the myocardium. DNA counterstaining was performed with DAPI (D) qPCR shows neural crest stem cell markers (*SOX10, SOX9, TWIST, SLUG*) next to markers for stem cells (*NESTIN*) and pluripotency (*MYC, OCT4*) in the heart auricle tissue.

### Successful isolation of putative human cardiac stem cells from heart auricle tissue

To analyse the NCSC-marker expressing cells found in the adult human myocardium in more detail, we modified an established protocol for the isolation of human cardiomyocyte progenitor cells ^4^. Explant culture resulted in the isolation of cells migrating out of tissue pieces (Fig. 2 A, Fig. S1 A+B) that spontaneously formed cardiospheres (Fig. 2 B).

**Figure 2:**
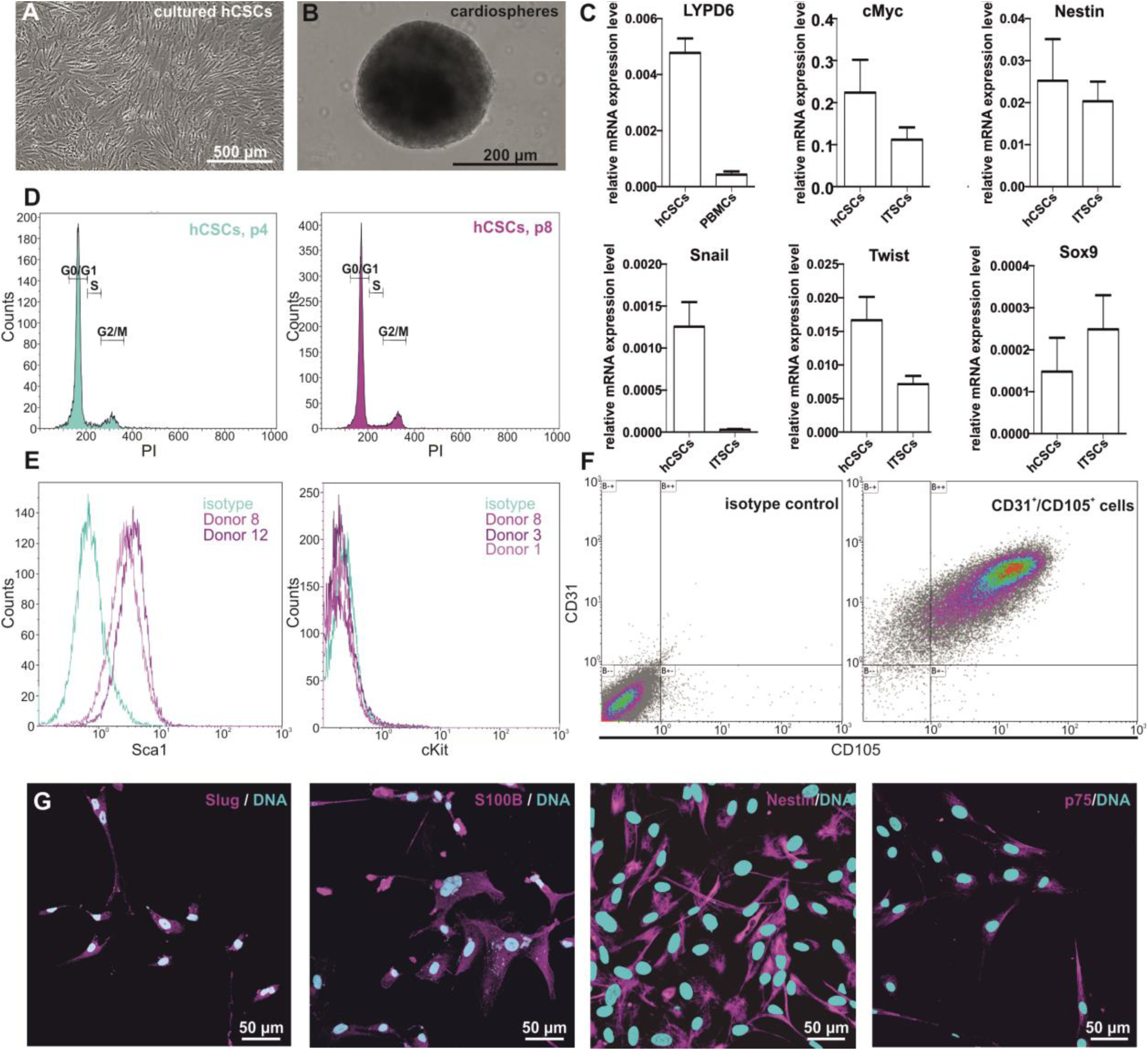
Successful isolation and in vitro expansion of hCSCs. (A) Adherent cell culture. (B) hCSCs spontaneously form cardiospheres during culture. (C) Cultured hCSCs express stem cell and pluripotency markers (*MYC, NESTIN*) as well as NCSC-markers (*SNAIL, TWIST, SOX9*) at comparable levels with known neural crest-derived stem cells (ITSCs). LY6-Family genes are expressed at comparable levels with PBMCs. (D) High passage-hCSCs show no aberrant DNA content compared to low passage-hCSCs. (E) cultured hCSCs express the cell surface marker Sca1, as detected by flow cytometry. cKit is not expressed. (F) Doublestaining shows that 92,56 % of cells are double positive for CD31 and CD105. Representative data of two independent donors. (G) Immunocytochemistry shows the expression of the neural crest stem cell markers Slug, S100B, Nestin and p75 in vitro. DNA counterstaining was performed with DAPI.

### Isolated putative human cardiac stem cells express neural crest markers on mRNA level

Analysis of the expression pattern showed the expression of the NCSC-markers *SNAIL, TWIST, SOX9* and *NESTIN* as well as the pluripotency marker *CMYC* in cell populations from 2 distinct donors (Fig. 2 C). Expression of NCSC-markers was observed to be at comparable level to adult human inferior turbinate stem cells (ITSCs), a well-characterized human neural crest-derived stem cell population we previously identified in the inferior turbinate of the nasal cavity ^15,18,24,31-33^.

### Putative hCSCs keep their genetic stability during cultivation in vitro

After long-term *in vitro* culture, a comparison of the DNA-content of PI-stained hCSCs in high and low passages revealed no changes in the PI signal above the G2/M peak and therefore no polyploidy in higher passages (Fig. 2 D). In summary, these findings indicate the successful isolation of neural crest-derived stem cells from adult human heart auricles.

### Sca1^+^/CD105^+^/CD31^+^/cKit^-^putative human cardiac stem cells are positive for neural crest markers Slug, S100B, Nestin and p75 on protein level

Characterizing putative hCSCs in more detail, we further observed a high amount of 87 % to 95.6 % of Sca1^+^ cells isolated from two distinct hCSC-donors but no expression of cKit using flow cytometry (Fig. 2 E). Multiple staining followed by flow cytometric measurement showed 92.56 % of cells being double positive for CD105 as well as CD31 (Fig. 2 F). Isolated and cultured putative hCSCs were also positive for the neural crest-specific markers Slug, S100B, Nestin and p75^NTR^, as detected by immunocytochemistry (Fig. 2 G).

### HCSCs show high clonal efficiency and capability for secondary cardiosphere formation

To investigate their stemness characteristics, the clonal growth of putative hCSCs was analysed (Fig. 3 A). Here, putative hCSCs revealed a clonal efficiency of 22.7 %. Importantly, clonally grown cells maintained their ability to form secondary cardiospheres under suspension culture conditions (Fig. 3 B).

**Figure 3:**
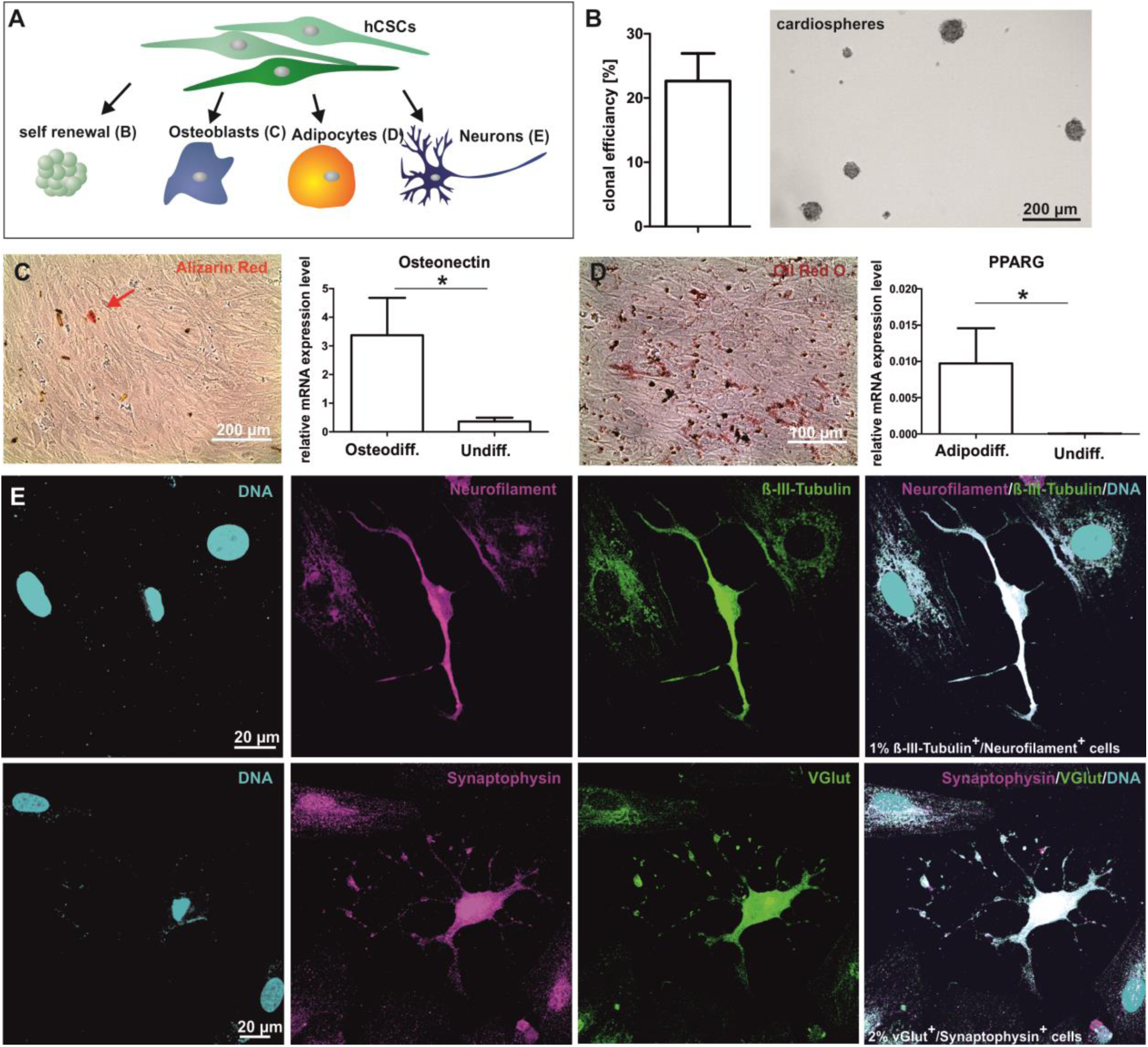
hCSCs differentiate into several cell types and are capable for self-renewal. (A) directed differentiation led to differentiation of hCSCs into ectodermal and mesodermal cell types. (B) hCSCs show a clonal efficiency of 22,7 % and are capable to form secondary cardiospheres. (C) Alizarin red staining detects calcium deposits of osteoblasts. qPCR shows that osteonectin is upregulated in comparision to undifferentiated cells. (D) Oil Red O staining detects hCSC-derived adipocytes. qPCR shows that peroxisome proliferator-activated receptor gamma (PPARG) is upregulated in hCSC-derived adipocytes compared to undifferentiated cells. (E) hCSCs differentiate into neuron-like cells expressing the neuronal markers Neurofilament, ß-III-Tubulin, Synaptophysin and VGlut. Nuclear counterstaining was performed with DAPI.

### HCSCs show the capability to differentiate into osteoblasts, adipocytes and neuron-like cells

Determining their capability to give rise to specialized cell types, we assessed the osteogenic, adipogenic and neuronal differentiation potential of hCSCs by applying established protocols (Fig. 3 A). HCSCs were able to differentiate into osteoblasts as indicated by alizarin red stained calcium deposition and increased expression of the specific osteogenic transcript Osteonectin in comparison to undifferentiated control (Fig. 3 C). Further, adipogenically differentiated cells could be detected by Oil Red O Staining and an upregulated gene expression of peroxisome proliferator-activated gamma (PPARG) compared to undifferentiated control (Fig. 3 D). As one example for the ectodermal differentiation potential of hCSCs, application of a defined neuronal differentiation protocol led to the formation of Neurofilament^+^/ß-III-Tubulin^+^ as well as Synaptophysin^+^/vGlut^+^ neuron-like cells (Fig. 3 E). Given the successful differentiation into derivates of both ectoderm and mesoderm, our findings indicate hCSCs to be a multipotent stem cell population with neural crest origin.

### HCSCs are able to give rise to beating cardiomyocytes in vitro

To investigate the cardiogenic differentiation potential of the hCSC-population, we exposed the cells to TGFβ and ascorbic acid as published by Smits and colleagues ^4^ (Fig. 4 A). A high amount of 94 % α-actinin^+^ cells was detected by immunocytochemistry, indicating a successful cardiac differentiation whereas no α-actinin was observable in undifferentiated control cells (Fig. 4 B+C). Additionally, these cells showed a characteristic striated pattern in the α-actinin immunostaining (Fig. 4 D). Despite the expression of marker proteins in nearly all of the differentiated cells, spontaneous beating was not observable after differentiation with TFGβ and ascorbic acid. To assess the capability of hCSCs to undergo differentiation into functional cardiomyocytes, we performed coculture experiments of GFP-transduced hCSCs and primary neonatal mouse cardiomyocytes (Fig. 4 A, E). After 11 days of coculture, we observed beating hCSC-derived GFP^+^ cardiomyocytes (Fig. 4 F, supplemental video), demonstrating successful cardiogenic differentiation of hCSCs.

**Figure 4:**
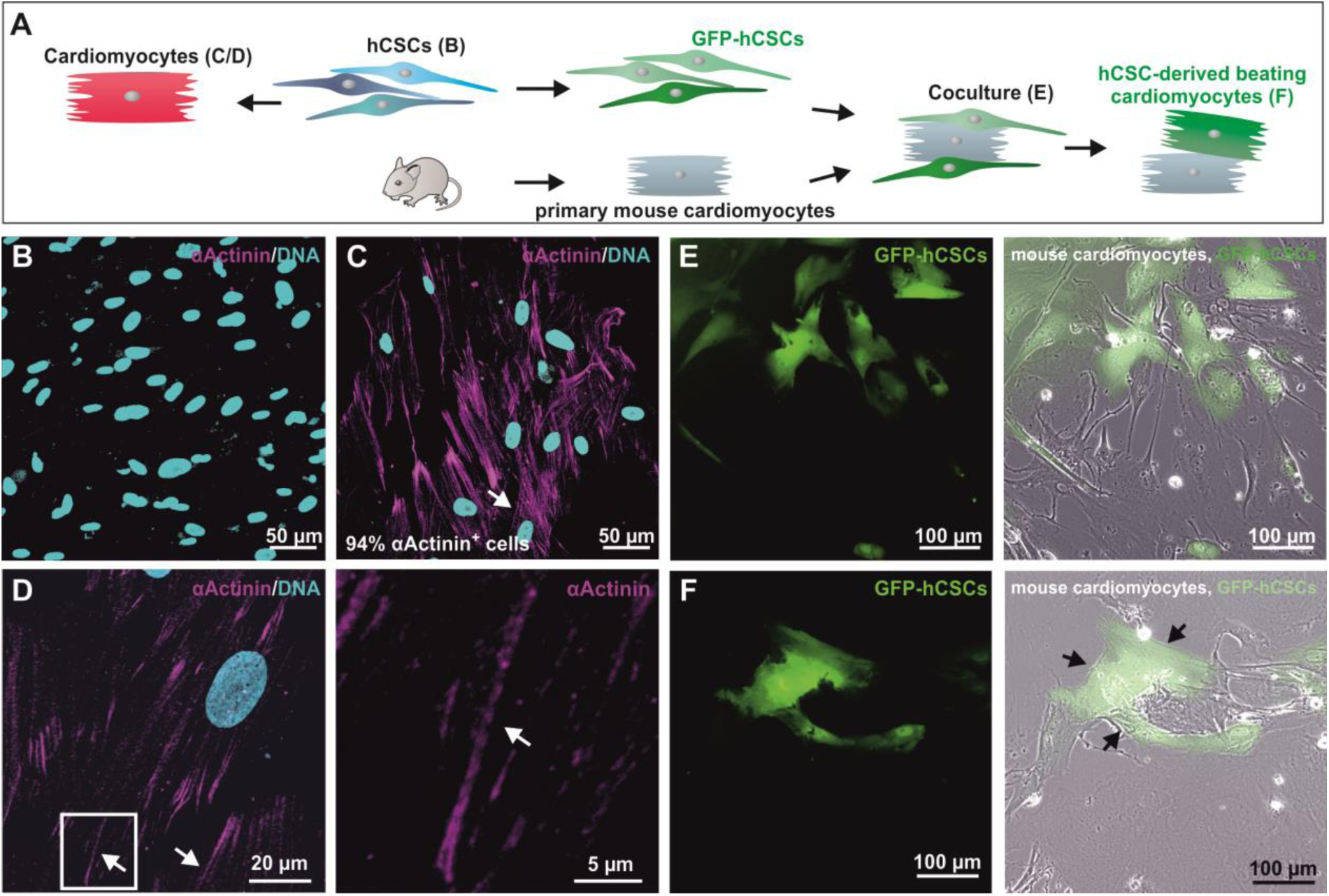
hCSCs differentiate into cardiomyocytes and beat spontaneously when cocultured with primary mouse cardiomyocytes. (A) Two different protocols were applied for the myocardial differentiation of hCSCs. (B+C) Application of TGFβ and ascorbic acid leads to differentiation of hCSCs into cardiomyocytes with 94 % αActinin^+^ cells compared to 0 % αActinin^+^ cells in undifferentiated controls. (D) αActinin-Immunostaining reveals a striated pattern in cardiogenic differentiated cells. Nuclear counterstaining was performed with DAPI. (E) Day 3 of coculture of Lentiviral transduced GFP^+^ hCSCs and primary neonatal mouse cardiomyocytes. (F) Day 12 of coculture of Lentiviral transduced GFP^+^ hCSCs and primary neonatal mouse cardiomyocytes. Arrowheads indicate beating GFP^+^ human cardiomyocytes (see also supplementary Movie M1).

### Global gene expression analysis show highly similar patterns between hCSCs and adult neural crest-derived stem cells despite donor dependent variations

To investigate the potential neural crest origin of hCSCs in more detail, we performed RNAseq of hCSCs from 4 distinct donors as well as of neural crest-derived inferior turbinate stem cells (ITSCs) from 4 different donors. In a previous study, we showed a great difference in gene expression between ITSCs and human embryonic stem cells^18^. In the present study, we found a high Pearson correlation between stem cell populations from different donors of the same tissue ranging from R^2^= 0.93 to 0.968 for ITSCs and R^2^=0.916 to 0.957 for hCSCs (Fig. 5 A). Surprisingly, the same range of correlation was measured between hCSCs and ITSCs independent to the respective donors with R^2^=0.898 to 0.929 (Fig. 5 B). A principal component analysis (PCA) showed that hCSCs and ITSCs form distinct clusters when compared with each other with PC1 explaining 30.57 % of the variance (Fig. 5 C) but both clusters show higher differences to populations of embryonal stem cells (ESCs) or induced pluripotent stem cells (iPSCs) where PC1 explains 62.04 % of the variance (Fig. 5 D) ^34^. Taken together, these data show a highly similar gene expression profile of hCSCs and ITSCs suggesting a similar developmental origin of both stem cell populations.

**Figure 5:**
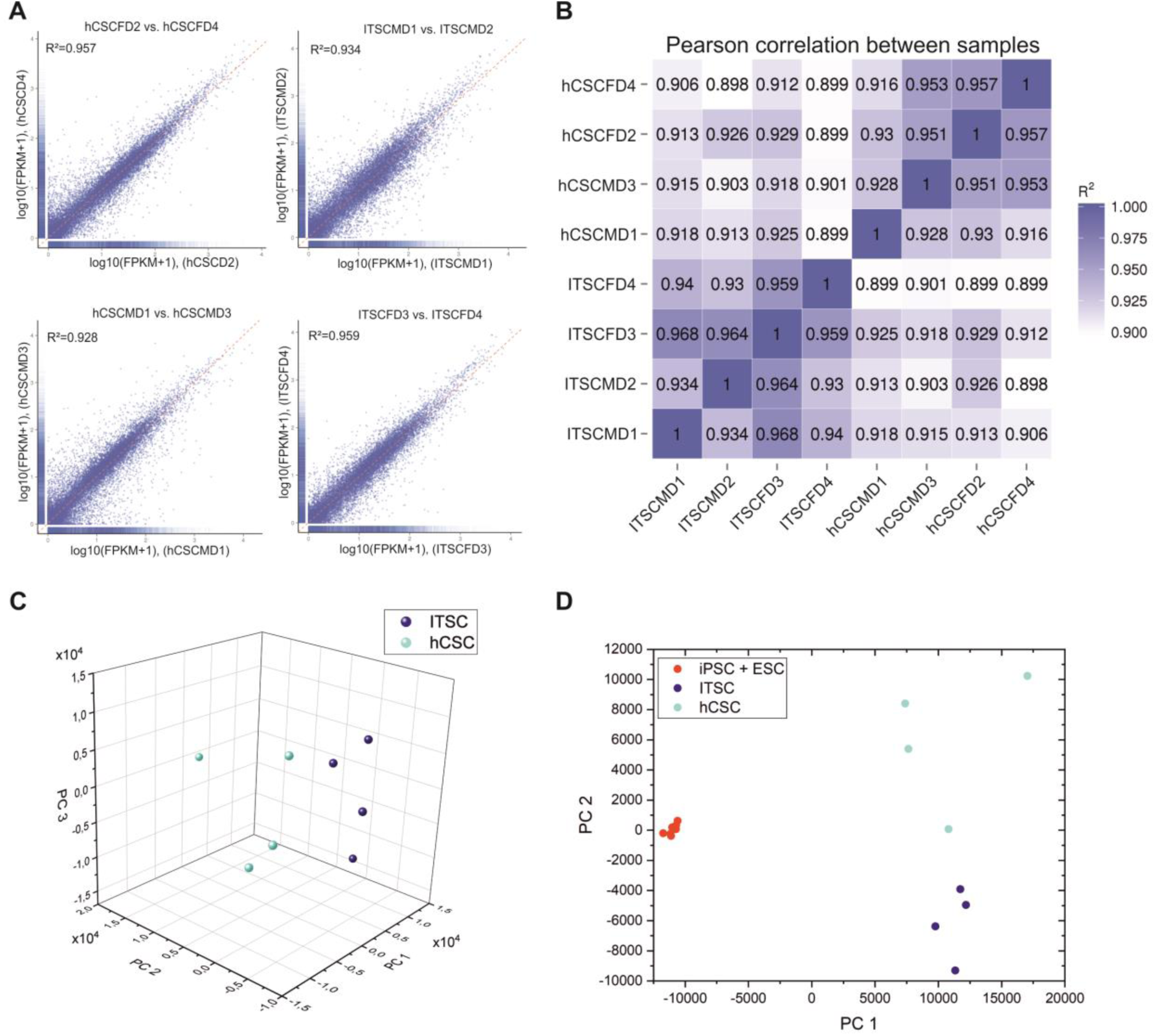
Global expression profiles of hCSCs and ITSCs correlate very high. (A) Scatter plots of two samples within a population show a correlation >92 %. (B) Pearsons’s correlation between samples of different populations averages 90 %. (C) Principal component analysis (PCA) shows that the gene expression patterns of hCSCs and ITSCs build distinct clusters (Significance: PC1 = 30,57 %, PC2 = 28,22 %, PC3 = 20,62 %). (D) PCA comparing hCSCs, ITSCs and ESCs and iPSCs (Significance: PC1 = 62,04 %, PC2 = 11,50 %).

We used different Venn diagrams to visualize similarities and differences of CSC and ITSC expression patterns in more detail. We detected 79 % of all expressed genes (10913 genes) to be similarly expressed in ITSCs of all 4 donors (Fig. 6 A). Furthermore, we did a similar analysis with hCSCs and detected an even higher number of genes being expressed in hCSCs of all 4 donors (11132 genes, 78 % of all expressed genes) (Fig. 6 B). Interestingly, across stem cell populations from different tissue types, the accumulated number of significantly expressed genes (11847, 88 %, Fig. 6 C) in cells cultivated from both tissue types was even higher than within one tissue type (see Fig. 5 C / Fig. 6 A, B for comparison). These data hind to a high similarity of hCSC- and ITSC expression profiles. Within this overlap of significantly expressed genes in both adult human stem cell populations, the stem cell markers *NESTIN* and *LYPD6* (a member of the Ly6-family like the mouse gene Sca1) could be found next to several NCSC-markers like *SNAIL, SLUG, TWIST* and *SOX9*, as well as the pluripotency genes *MYC* and *KLF4* (Fig. 6 D). Interestingly, cKit and Oct4 expression could not be detected in both stem cell populations. Taken together, known NCSC markers are expressed in both stem cell populations, further strengthening the hypothesis that hCSC are of neural crest origin.

**Figure 6:**
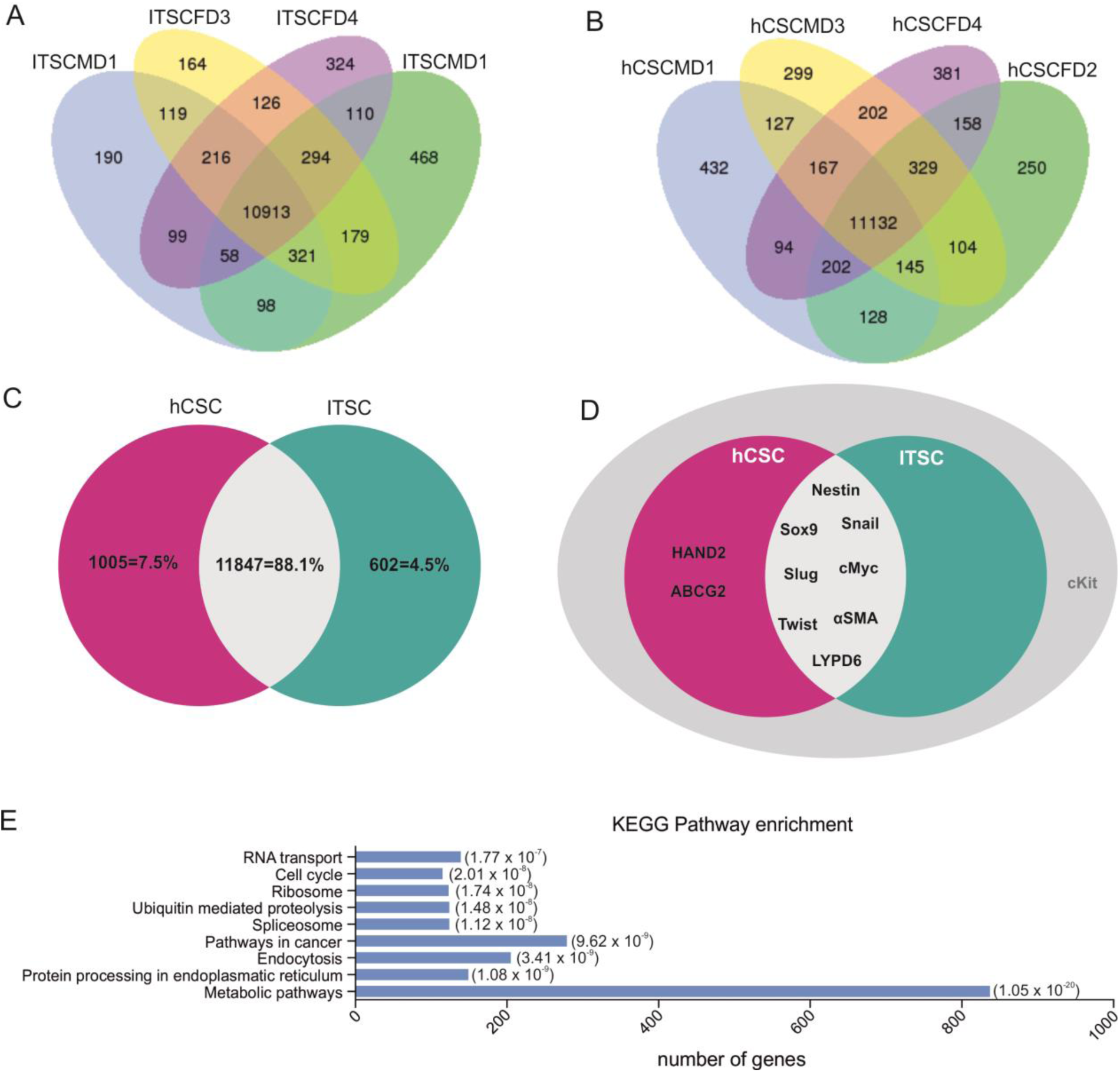
hCSCs and ITSCs coexpress a wide range of genes. (A) Venn diagramm of coexpressed genes within all analysed ITSC donors. (B) Venn diagramm of coexpressed genes within all analyzed hCSC donors. (C) Overlap of significantly expressed genes by hCSCs and ITSCs is 88,1%. (D) Coexpressed genes include Nestin, Sox9, Snail, Slug, Twist, cMyc, Sca1 and αSMA and exclude Lgr5, Oct4 and c-kit. (E) KEGG Pathway enrichment of similar expressed genes. Top 9 most significantly enriched KEGG pathways are inidcated.

### KEGG Pathway and Gene Ontology Analysis show a variety of mitosis-associated terms significantly enriched in hCSCs and ITSCs

Investigation of similar expressed genes in hCSCs and ITSCs using KEGG Pathway enrichment led to the discovery of 128 significantly enriched KEGG Pathway terms. Among the 10 most significant enriched pathways, we found the terms metabolic pathways (ID: hsa01100, corrected p-value: 3.13 × 10^−20^), protein processing in endoplasmatic reticulum (hsa04141, 1.08 x 10^−9^), cell cycle (hsa04110, 2.01 × 10^−8^) and RNA transport (hsa03013, 1.77 × 10^−7^) (Fig. 6 E). We also found the stem cell-associated KEGG-terms hippo signalling pathway (hsa04390) and signalling pathways regulating pluripotency of stem cells (hsa04550) (data not shown). To further reduce the RNAseq data dimensionality, we performed Gene Ontology (GO) analysis of the similar expressed genes in ITSCs and hCSCs. Here, we detected among the top 20 biological processes of the most enriched GO-terms a variety of mitosis-associated terms like “signal transduction involved in cell cycle check point” (GO:0072395), “signal transduction involved in DNA integrity checkpoint” (GO:0072401) or “signal transduction involved in DNA damage checkpoint” (GO:0072422) (Fig. 7 A). In addition, biological processes like “cardiovascular system development” (GO: 0072358), “nervous system development” (GO: 0007399), “developmental processes” (GO: 0032502), “neuron fate specification” (GO: 0048665) and “regulation of multicellular organismal development” (GO: 2000026) could be found among the enriched GO terms (data not shown). Accordingly, among the top 20 most enriched GO-terms of cellular components, we found nucleus associated terms like “condensed chromosome centromeric region” (GO:0000779), “condensed chromosome kinetochore” (GO:0000777) and “spliceosomal complex” (GO:0005681) (Fig. 7 B). Moreover, enrichment of the category molecular function revealed GO-terms like “RNA binding” (GO:0003723), “single stranded DNA binding” (GO:0003697) and “helicase activity” (GO:0004386) (Fig. 7 C). These results strongly suggest a similar regulation of cell cycle and mitosis in hCSCs and ITSCs.

**Figure 7:**
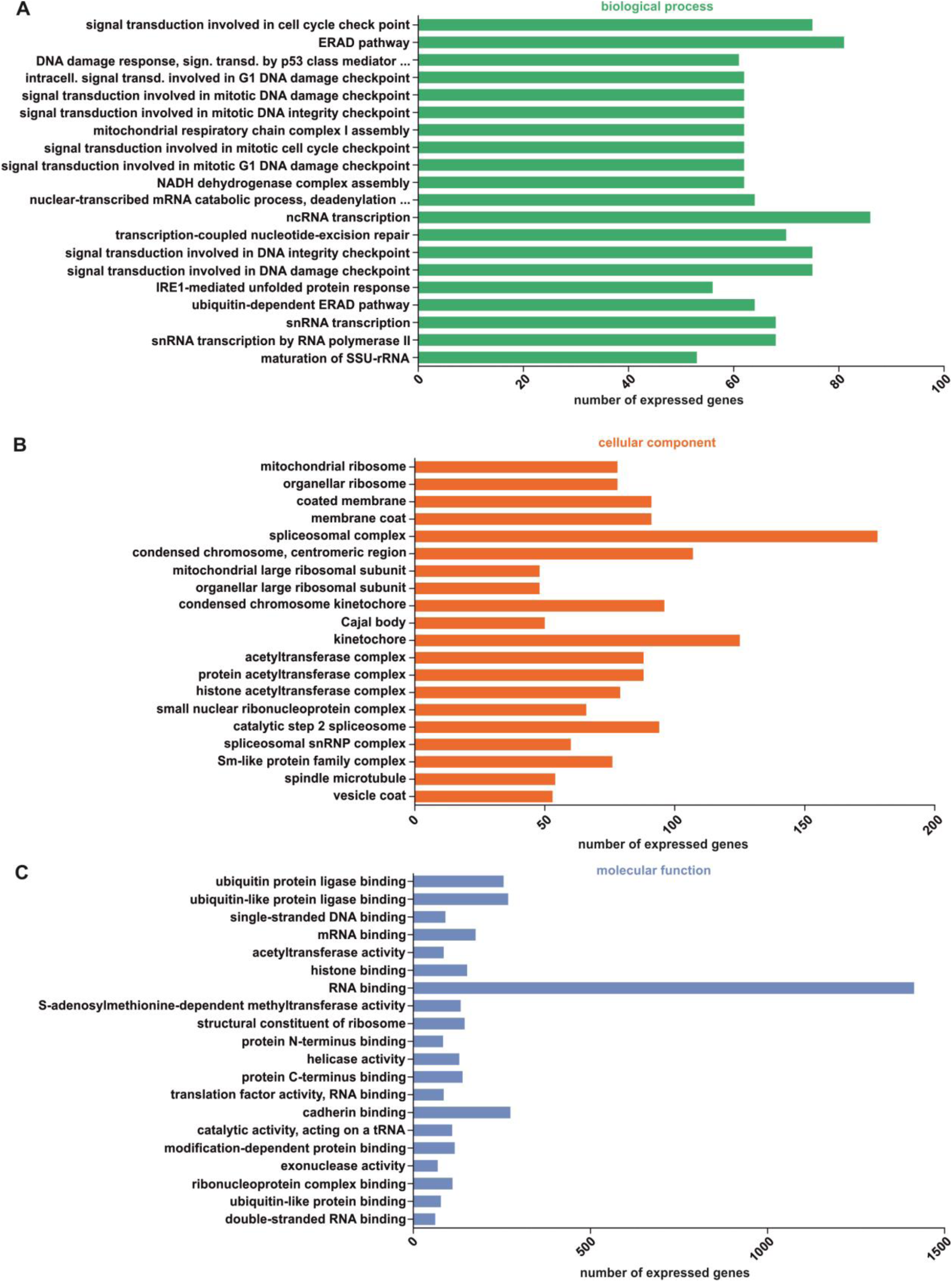
GO-term enrichment of similar expressed genes in hCSCs and ITSCs. (A) Top 20 enriched GO terms of biological processes. (B) Top 20 GO terms associated with cellular component. (C) Top 20 GO terms associated with molecular function.

## DISCUSSION

The present study describes a Nestin^+^/S100B^+^/CD105^+^/Sca1^+^/cKit^-^ population of human cardiac stem cells in the myocardium of the adult human heart auricle. These cells express several NCSC and pluripotency markers in their niche as well as during cultivation in vitro and give rise to beating cardiomyocytes, osteoblasts, adipocytes and neurons. Further, by a global transcriptome analysis via RNAseq, we were able to show high correlation in the expression profile of Nestin^+^/S100B^+^/CD105^+^/Sca1^+^/cKit^-^ hCSCs with a human NCSC population from the nasal cavity, demonstrating the neural crest-origin of this human cardiac stem cell population. Although a range of stem cell populations was described to be present in the human heart^3,4,35^, a potential relation of these human cardiac stem cell populations to the neural crest has not been described so far. The neural crest was initially described by Wilhelm His in the development of the chick embryo as the intermediate chord appearing between the neural chord and the future ectoderm ^14^. After neurulation, neural crest cells migrate to a broad range of target tissues within the developing organism and give rise to various cell populations cells of mesodermal and ectodermal type ^32^. In addition, neural crest-derived cells also persist as adult stem cell populations within the adult body^18-21,36^. Extending the range of known NCSC-populations within the human organism, our present findings show the existence of an adult NCSC-population within the myocardium of the adult human heart auricle. Accordingly, a potential developmental relation of cardiac stem cells to the neural crest has already been suggested in mice and rats^22,23,37,38^. In particular, El-Helou and colleagues demonstrated the presence of NCSCs in the adult rat heart via expression of Nestin ^37^, a characteristic NCSC-marker associated with proper self-renewal of stem cells ^32,39^, while Tomita and coworkers could show, that Nestin^+^ NCSCs in the mouse heart give rise to cardiomyocytes *in vivo* ^40^. In the present study, we likewise identified Nestin-positive cells in the human heart, which expressed a range of neural crest and stemness markers including *SNAIL, SLUG, SOX9, TWIST*, and *cMYC*. Notably, similar to NCSCs populations residing within the head and neck region ^15^-^19^, Nestin^+^ human cardiac NCSCs exhibited an extraordinary high differentiation capability by giving rise to cardiomyocytes but also other mesodermal and ectodermal cell types. These observations extend the commonly described differentiation ability of human CSCs, which was reported to be restricted to cardiomyocytes, smooth muscle cells and endothelial cells *in vitro*^4,5^. This extended multipotency of the here presented adult hCSC-population is described as a characteristic hallmark of neural crest-derived stem cells ^16^, making NCSCs to highly promising candidates in tissue repair and regenerative medicine ^15,31,41^.

Identification and isolation of adult CSCs is commonly focused on the expression of the cell-surface marker cKit^2,5,42^. The cKit receptor is a type III tyrosine-protein kinase that binds the stem cell factor (SCF) as ligand. Its expression has also been shown to be important for the proliferation of hematopoietic stem cells (HSCs) and is used as a marker for HSCs^43^. cKit is also expressed in interstitial cajal cells of the human embryo^44,45^, indicating the expression of the cKit protein to be not cardiac-specific and therefore not sufficient to identify cardiac stem cells. In addition, the relevance of cKit as a cardiac stem cell marker is controversially discussed. For instance, Sultana and co-workers showed, that cKit^+^ cells in the mouse heart are endothelial cells and not cardiac stem cells^46^. Unfortunately, such studies do not exist for the human system. Accordingly, we show that the myocardium of the adult human heart auricle contains a population of Nestin^+^/S100B^+^ cardiac stem cells lacking cKit-expression in the present study. Next to cKit, the surface antigens Sca1 and CD105 are commonly used markers for human cardiac stem and progenitor cells ^4^. In contrast to most studies showing co-expression of Sca1, cKit and CD105 in human CSCs ^4^, we observed the presence of Sca1 and CD105 in human cardiac NCSCs despite the lack of cKit.

As already postulated by Iancu and colleagues, it needs not only the presence of single cell surface markers but rather an extensive marker profile, consisting of a combination of specific cell surface markers and global gene expression to distinguish distinct cardiac and non-cardiac stem cell populations ^47^. In this regard, we performed RNAseq to compare global transcriptional profiles of isolated CSCs from the human heart auricle with NCSCs from the human nasal cavity, namely inferior turbinate stem cells (ITSCs)^18^. Pearson’s correlation coefficients of pairwise comparisons exceeded 92% within both populations, indicating high similarities in the respective transcriptional profiles and thus an affiliation to a distinct homogeneous population. For comparison, Vicinanza and co-workers accepted a Pearson’s correlation of 81% between two samples of the same CSC-population as sufficient homogenous population^48^. Further, R^2^ between samples of Nestin^+^ hCSCs and ITSCs populations were approximately 90% for all combinations. In addition, principal component analysis (PCA) showed a distinct cluster including ITSCs and hCSCs, which was clearly separated from iPSCs and ESCs and underlined the high similarity between ITSCs and hCSCs in terms of gene expression profiles. Transcriptome analysis of ITSCs and hCSCs revealed an overlap of 88.1 % genes that were significantly expressed in both cell populations, including the neural crest markers *NESTIN, SNAIL, SLUG, TWIST* and *SOX9*. In terms of gene expression profiling, expression of *HAND2*, a transcription factor playing an essential role in cardiac development^49^, was unique in hCSCs.

Next to identifying hCSCs as a novel NCSC-population, our findings likewise show a surprisingly high correlation in global gene expression between human NCSCs from the trunk region (heart) and the cranial region (nasal cavity). Although NCSC-populations from cranial and trunk region were not compared using global gene expression profiles so far, commonly accepted differences were described for mice und chicken in terms of their differentiation potential ^50^. Accordingly, we observed differences in the differentiation potential of both stem cell populations particularly regarding the extraordinary high differentiation capability of ITSCs into the neuronal lineage^15^. Despite this differences in their developmental potential, which we suggest to depend on the particular niche of the stem cell populations, our finding emphasize a strong similarity between NCSCs from the cranial and trunk region in the human organism.

In summary, we describe the presence of a Nestin^+^/S100B^+^/CD105^+^/Sca1^+^/cKit^-^ population of adult neural crest-derived stem cells within the adult human heart. The here identified Nestin^+^/S100B^+^/CD105^+^/Sca1^+^/cKit^-^ hCSC-population may contribute to endogenous cardiac tissue homeostasis and tissue repair *in vivo* and may serve as promising cell source for future therapeutic applications.

## ACKNOWLEDGEMENTS

This work was supported by the Fund for the promotion of transdisciplinary, medically relevant research cooperations in the region Ostwestfalen-Lippe as well as by the University of Bielefeld and the Heart and Diabetes Centre NRW.

## AUTHOR CONTRIBUTIONS STATEMENT

A.H. Collection and assembly of data, data analysis and interpretation, manuscript writing, final approval of manuscript. M.M., K.S. and L.R. Collection and assembly of data, data analysis and interpretation, final approval of manuscript. I.F., J.Gr. Data analysis and interpretation, manuscript writing, final approval of manuscript. H.H., T.H. Data analysis and interpretation, final approval of manuscript. S.S., B.F. and T.P. and J.Gu. Collection and assembly of data, final approval of manuscript. B.K., C.Kn. and C.Ka. Conception and design, financial support, data analysis and interpretation, manuscript writing, final approval of manuscript.

---

**Supplemental Figure S1:**
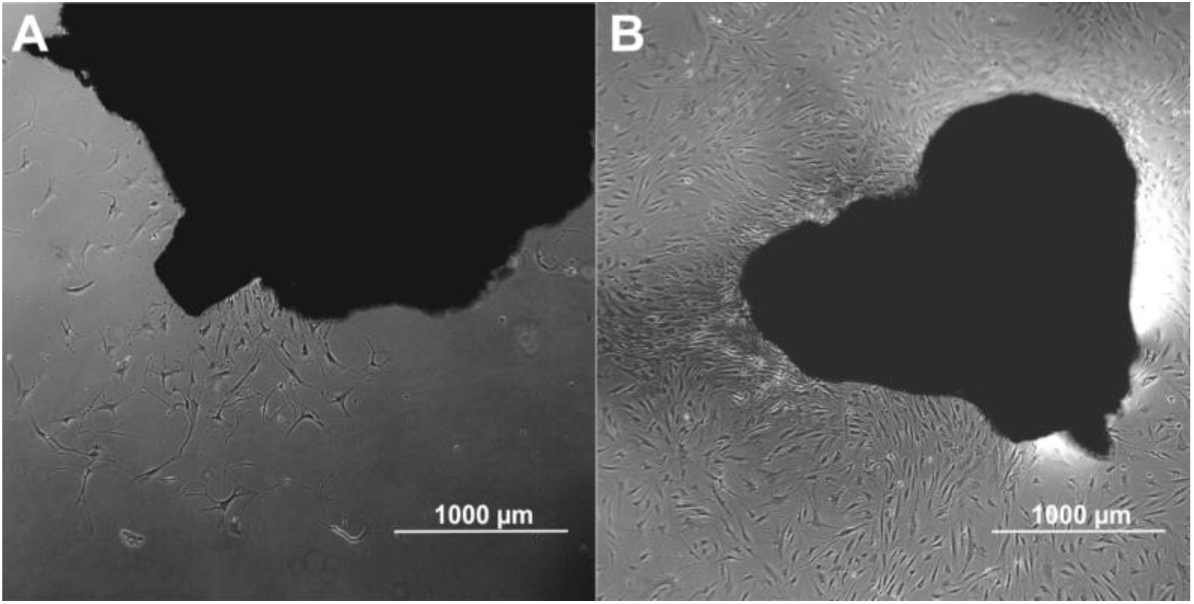
Explant culture of human heart auricles. Tissue clumps are placed on gelatine B-coated tissue culture flasks. (A) Cells start to migrate out of the tissue at day 7 of explant culture. (B) A confluent monolayer of cells can be observed at day 14 of explant culture.

